# Accumulation of somatic mutations leads to genetic mosaicism in Cannabis

**DOI:** 10.1101/2021.02.11.430823

**Authors:** Kristian Adamek, Andrew Maxwell Phineas Jones, Davoud Torkamaneh

## Abstract

Cannabis is typically propagated using stem cuttings taken from mother plants to produce genetically uniform propagules. However, producers anecdotally report that clonal lines deteriorate over time and eventually produce clones with less vigour and lower cannabinoid levels than the original mother plant. While the cause of this deterioration has not been investigated, one potential contributor is the accumulation of somatic mutations within the plant. To test this, we used deep sequencing of whole genomes (>50x) to compare the variability within an individual *Cannabis sativa* cv. “Honey Banana” plant sampled at the bottom, middle and top. We called over 6 million sequence variants based on a reference genome and found that the top had the most by a sizable amount. Comparing the variants among the samples uncovered that nearly 600K (34%) were unique to the top while the bottom only contained 148K (12%) and middle with 77K (9%) unique variants. Bioinformatics tools were used to identify mutations in critical cannabinoid/terpene biosynthesis pathways. While none were identified as high impact, four genes contained more than double the average level of nucleotide diversity (π) in or near the gene. Two genes code for essential enzymes required for the cannabinoid pathway while the other two are in the terpene pathways, demonstrating that mutations were accumulating within these pathways and could influence their function. Overall, a measurable number of intra-plant genetic diversity was discovered that could impact long-term genetic fidelity of clonal lines and potentially contribute to the observed decline in vigour and cannabinoid content.

## Introduction

*Cannabis sativa* L. (marijuana, hemp, cannabis; Cannabaceae) is regarded as one of the first crops humans domesticated and is primary a dioecious diploid annual species (2*n* =20) cultivated for fiber, oil, seed and its medicinal and psychoactive properties (Hillig, 2005). The main pharmaceutical and psychoactive compounds are cannabinoids that accumulate in trichomes produced primarily on floral tissues of female plants (van Bakel *et al*., 2011). To date, 177 cannabinoids have been identified and described, with the two most naturally abundant, well-studied, and sought after being (–)-*trans*-Δ^9^–tetrahydrocannabinol (THC) and cannabidiol (CBD) (Hanuš and Hod, 2020; Hurgobin *et al*., 2020). These compounds have many demonstrated therapeutic properties, including alleviating symptoms in epilepsy (Devinsky *et al*., 2014) and multiple sclerosis (van Amerongen *et al*., 2018) and are being investigated for other ailments such as Alzheimer’s disease (Watt and Karl, 2017). Additionally, cannabis produces hundreds of different terpenes/terpenoids, which are also known to have therapeutic effects, including antifungal, antiviral, anticancer, anti-inflammatory, antiparasitic, antioxidant, and antimicrobial activities, and are thought to interact with cannabinoids to alter their activities (Hanuš and Hod, 2020). Cannabinoids, terpenes and other secondary metabolites are produced in the capitate stalked glandular trichomes and minor variations in concentrations of each may result in distinct therapeutic effects (Hurgobin *et al*., 2020).

The polyketide, cannabinoid, and methylerythritol phosphate (MEP) pathways are responsible for the creation of the cannabinoids (i.e., THCA, CBDA, CBCA, CBGA) and the monoterpene, mevalonate (MEV), sesquiterpene and methylerythritol phosphate (MEP) pathways produce several different terpenes. The medicinal effects of Cannabis plants depend on the relative concentration of these compounds and are often classified into three main categories based on the THC:CBD ratio. Type I plants express a well over 1:1 THC:CBD ratio, Type II plants have moderate amounts with a near equal ratio, and Type III plants contain a less than 1:1 ratio (Hurgobin *et al*., 2020). However, it should be noted that this is an oversimplification of the chemical diversity found within Cannabis and that each plant has a unique chemical fingerprint that may impact its biological activity.

Clonal propagation is the primary method used when cultivating Type I & II cannabis plants for medicinal or recreational use to ensure genetic and phenotypic uniformity (McKernan *et al*., 2020). To achieve this, mother plants are established from elite seedlings that have been selected largely based on their specific chemical profile. Mother plants are capable of supplying hundreds or thousands of cuttings, usually taken from the apical region of the plant. The mother plants are maintained in an indefinite vegetative state for many years using a constant long day photoperiod (18:6 hours) and are occasionally replaced using a clonal propagule taken from themselves.

Vegetative propagation is used in many other domesticated plants to maintain valuable genotypes including bananas, potatoes, grapes, hops, and coffee trees (McKey *et al*., 2010; Carrier *et al*., 2012). Theoretically, clones produce plants that are genetically identical and phenotypically similar to the parent stock, however, cannabis producers have observed a decline in the quality of clones taken from a mother plant, usually resulting in reduced cannabinoid production and plant vigour (Cannabis growers, personal communication, 2020). While no peer-reviewed study has investigated this in cannabis, it has been discussed widely in the gray literature (Burnstein, 2019) and this is a well-known phenomenon in other species that demonstrate a decline in plant vigour during extended periods of vegetative propagation (Muller, 1932; Muller, 1964).

Muller’s Rachet, a term first proposed by Felsenstein (1974), is a theory developed to explain this phenomenon and suggests that in the absence of sexual recombination, species accumulate irreversible somatic mutations (Muller, 1932; Govindaraju *et al*., 2020). Further, since the majority of random mutations are deleterious, the long-term effect of their accumulation is a decline in plant vigour similar to what has been observed in cannabis. While somatic mutations have also positively impacted agriculture by producing unique clonal varieties of apples, citrus, and wine grapes derived by propagating genetically diverse bud sports, the phenomenon is problematic for long term genetic preservation of elite individuals. Whether somatic mutations are beneficial or deleterious, it is well-known that they occur and can lead to genetic diversity even within a single plant.

The source of this diversity lies in the nature of plant growth and development, in which meristematic regions grow and develop independent from one another. As they grow, they each accumulate a unique set of somatic mutations that leads to genetic diversity within a plant. This phenomenon has been under investigation since the 1970s where isozyme analysis was initially used to demonstrate genetic variation within single trees (Marshall and Allard, 1970). More recently, DNA sequencing verified this and identified that genetic distance increases systematically throughout a tree such that the mutation load is greatest at the distal end (Diwan *et al*., 2014; Orr *et al*., 2020). This phenomenon is known as the Genetic Mosaicism Hypothesis (GMH) and states that individual plants become genetically diverse due to the accumulation of spontaneous mutations occurring randomly as independent branches grow (Gill *et al*., 1995). While this phenomenon is often neglected during preservation of clonal lines, it could have a significant impact on long term genetic fidelity, especially in species with higher-than-average mutation rates.

In this study, we examined if the GMH applies to cannabis by sequencing the full genome of three samples taken from different locations within a single mother plant. These data were used to identify nucleotide variants within the plant using bioinformatics tools and assess the degree of variation within the plant. We then calculated the unique and shared variants among samples. Lastly, we investigated the potential impact of the nucleotide variants in critical cannabinoid and terpene synthesis genes. Overall, this study confirms there was a significant degree of genetic variability within a single cannabis plant and raises concerns about long term genetic preservation using clonal propagation.

## Results

### Deep full genome sequencing of bottom, middle and top regions from a mother plant

We performed deep whole-genome sequencing (WGS) on three samples, taken from the stems located at 59 cm, 151 cm, and 226 cm from the top of the pot of a 2.4 m tall, 1.5-year-old mother plant (Figure 1A). This mother plant was a clonally propagated seedling selection from *Cannabis sativa* cv. “Honey Banana” (HB) plant (BrantMed Inc., Brantford, ON), which is a high THC type-I plant. In total, we generated >1 billion 150-bp paired-end reads using an Illumina Novaseq 6000 technology (Table 1). This represents an average 58x depth of coverage across the three samples. These reads were mapped against the public cannabis reference genome (cs10 v. 2.0; GenBank Accession No. GCA_900626175.2) with a mapping success rate of greater than 93% (Table 1), thus covering >97% of the cs10 v. 2.0 genome sequence (Grassa *et al*., 2021). Using the Fast-WGS (Torkamaneh *et al*., 2018) bioinformatics pipeline, we detected 1.3, 0.9, and 1.7 M nucleotide variants (i.e. single- and multiple nucleotide variants (SNVs and MNVs) and small insertions/deletions (indels)) in the samples derived from the bottom, middle, and top of the mother plant, respectively. Originally, 6.4 M nucleotide variants were detected but due to the high level of heterozygosity discovered, we employed a filter to both the minimum number of reads (minNR) and the minimum number of reads containing variants (minNV) to greater than 10. Altogether, over 3.8 M variants compared to the reference genome were identified with a transition/transversion (Ts/Tv) ratio of 1.9. As detailed in Table 1, the top of the plant had the most variants, followed by the bottom, while the middle had the fewest, demonstrating intra-plant genetic diversity. When comparing the locations there was a difference of ~400K variants between the bottom and middle as well as the bottom and top. However, the difference between the middle and top was much greater with a total of 850K variants.

**Figure 1.**
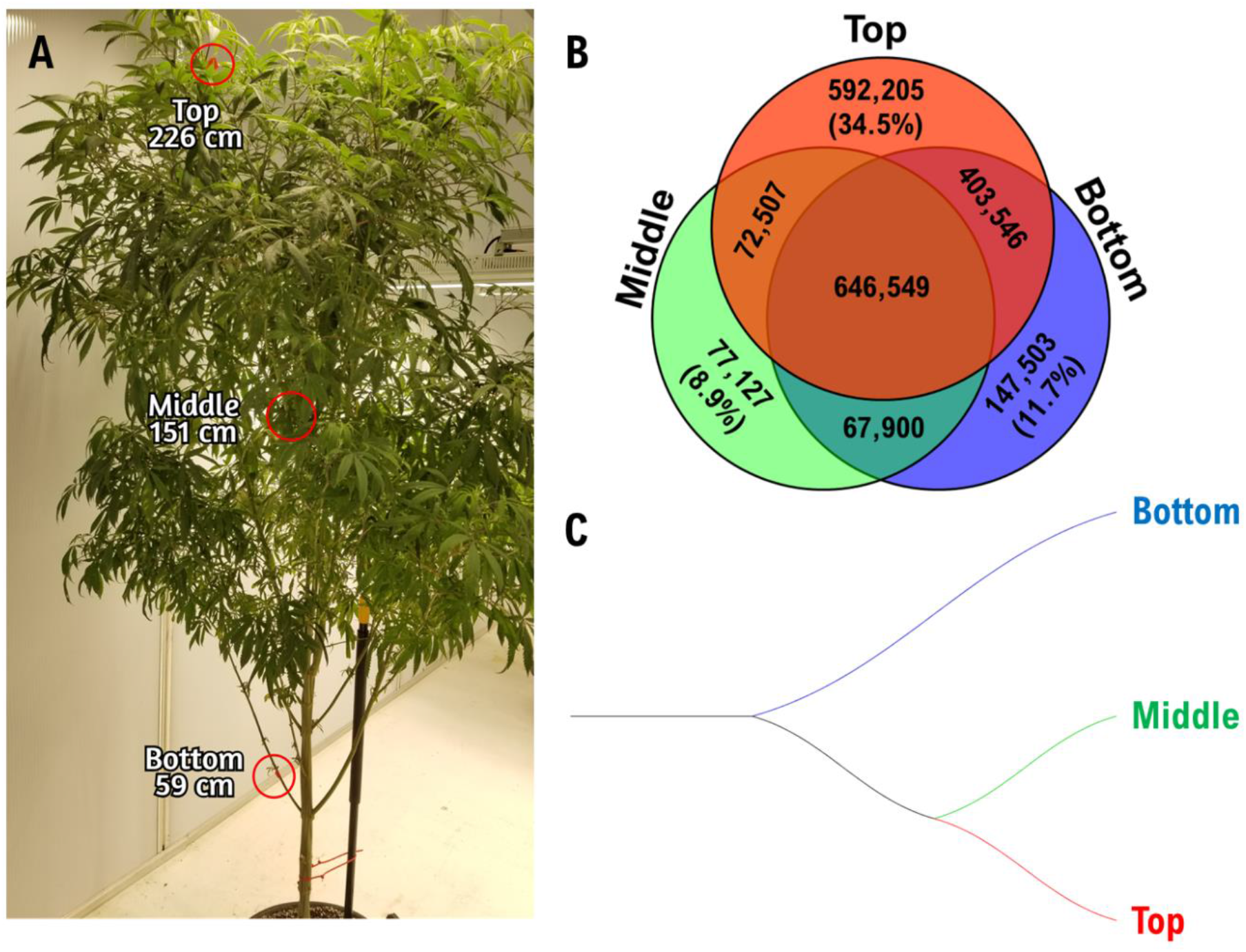
Cannabis mother plant with bioinformatic analysis. (**A**) Honey Banana, a mother plant (age: ~1.5 years), exhibited incredible growth, vigour and excellent traits which made this an ideal cultivar for this study. Three stem samples were taken from the top (226cm), middle (151cm) and bottom (59cm) and had their genomes fully sequenced. (**B**) A Venn diagram reveals the overlapping and unique nucleotide variants for the top, middle and bottom. (**C**) A phylogenic tree created using a genetic distance approach and the neighbour-joining method demonstrates that the top and middle are more similar by sharing a common node while the bottom has a separate branch.

**Table 1.**
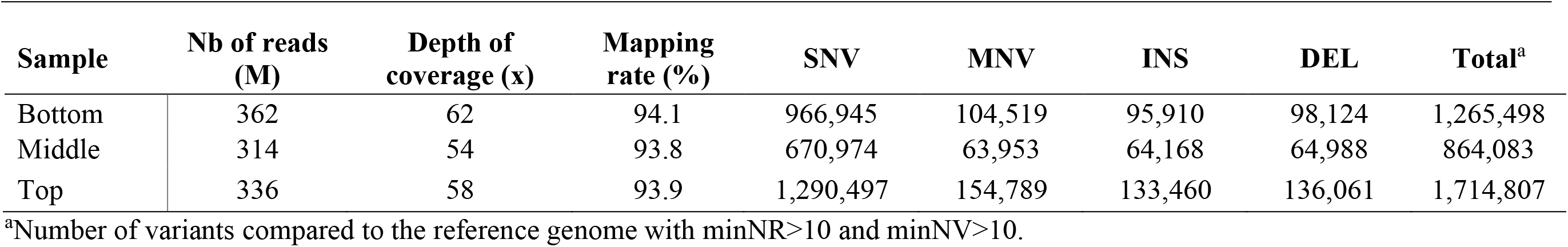
Statistics related to the whole-genome sequencing of Cannabis samples.

| Sample | Nb of reads<br>(M) | Depth of<br>coverage (x) | Mapping<br>rate (%) | SNV | MNV | INS | DEL | Total<sup>a</sup> |
| --- | --- | --- | --- | --- | --- | --- | --- | --- |
| Bottom | 362 | 62 | 94.1 | 966,945 | 104,519 | 95,910 | 98,124 | 1,265,498 |
| Middle | 314 | 54 | 93.8 | 670,974 | 63,953 | 64,168 | 64,988 | 864,083 |
| Top | 336 | 58 | 93.9 | 1,290,497 | 154,789 | 133,460 | 136,061 | 1,714,807 |

<sup>a</sup>Number of variants compared to the reference genome with minNR>10 and minNV>10.

### Variant calling quality assessment

To assess the quality of genotype calls, a second run of WGS was conducted on a new DNA library extracted, using similar protocols and procedures, from the original tissue from the top. In total, we generated 172 million 150-bp paired-end reads using an Illumina Novaseq 6000 technology representing an average of 32x depth of coverage across the cannabis genome. Using the Fast-WGS (Torkamaneh *et al*., 2018) bioinformatics pipeline and applying a set of supervised filters to exclude low-quality variant calls (details in M&M), we identified 1.7 M nucleotide variants with a 99.97% agreement in variant calling with the first experiment (Figure S1). This analysis suggests that an extensive and highly accurate set of nucleotide variants were obtained in this study.

### Mutational variation detected inside a singular *C. sativa* mother plant

For further analysis, the catalogues of detected variants relative to the cannabis reference genome (cs10 v. 2.0; (Grassa *et al*., 2021)) for each sample (i.e. bottom, middle, and top) were compared to assess the overlapping and unique variants. As can be seen in Figure 1B, more than 600K variants, compared to reference genome, were shared among all samples representing 51%, 75%, and 38% of variants detected in the sample from the bottom, middle, and top section of the mother plant, respectively. Bottom and top samples shared the greatest number of variants (403K), however top and middle shared only 72K variants. The lowest overlap was observed between bottom and middle, 67K. Most interestingly, the top sample contained the most unique variants at 34% (592K) followed by the bottom with 12% (147K) and the middle with 9% (77K). The most intriguing aspect of the result is that the top sample contained the most *de novo* (new) mutations. We document a very high rate of somatic mutations among the bottom, middle and top of a cannabis mother plant during vegetative growth.

According to the π statistic, the nucleotide diversity within three samples from a single plant was π = 6.0 × 10^−4^. To explore the genetic similarity between samples using nucleotide variants, we constructed a phylogenic tree using a genetic distance approach and the neighbour-joining method with repetition 1000x bootstrap test (Figure 1C). This shows two main branches, an individual bottom branch as well as a middle and top branch as the second, since they share a node. Within a single plant, we observed a very quickly declined linkage disequilibrium (LD) (Figure S2). The LD was seen to decay to its half in only a few kb.

Finally, based on a visualization approach using IGV (Robinson *et al*., 2011), we determined if the variants were clustered in specific regions of the genome. As can be seen from Supplemental materials Figure 3, there were certain areas in the genome that contain fewer mutations and others where elevated levels of mutations emerged. Specifically, one, five, eight and the X chromosomes had an apparent increased count while chromosomes two, three, four, six, seven and nine showed to have fewer. Furthermore, an intriguing observation was that chromosomes with more mutations also had higher levels of mutations in euchromatin regions (i.e. gene rich). Altogether, clusters of mutations appeared across the genome and as a result could indicate mutational hotspots exist within the cannabis genome.

### Functional impact on the genome from mutations are divergent depending on the location on the plant

To explore the potential functional impact of the mutations, we classified sequence variants into five categories based on their localization and identified the putative impact of the mutations. As can be seen in Figure 2A, more than half (51%) of the variants were in up/downstream regions, hence in close proximity of genes (5kb before and after gene) and the other 49% of the variants were located in intergenic regions (35%), genic regions (13%), splice sites (0.4%), and untranslated regions (UTR; 0.4%). Additionally, the genic category consists of exons, introns, and transcriptional variants at 2%, 4.5%, and 6.7%, respectively. From a functional standpoint, we were particularly interested in the subset of mutations predicted to have a large impact. Therefore, we explored the category of the high impact mutations (i.e., variants which are predicted to have a disruptive impact on the protein, probably leading to protein truncation, loss of function or triggering non-sense-mediated decay). Figure 2B presents the unique and shared high impact mutations from the bottom, middle and top samples. This information follows a similar pattern as seen for entire variants (Figure 1B) where the percentage of variants are similar. The number of shared high impact mutations between all samples was 845 which corresponds to 41%, 77%, and 28% of the total high impact mutations for the bottom, middle and top, accordingly. The top sample had the most unique high impact mutations with 1,234 (40%), next the bottom with 247 (12%) and lastly the middle with 63 (6%). The most intriguing was middle and top because there was a large difference between the unique variants (1.2K) (Figure 2B). Also, they shared 952 with 107 exclusively together, which represents 31% of top and 87% of middle total high impact mutations. To provide a more relevant perspective, we mainly focused on the high impact mutations from the middle and top samples as they are new mutations and ultimately, they might show a different phenotype compared to the original plant (i.e., bottom). In total, 1,333 high impact mutations were divided into four categories and represented frameshift (46%), premature stop codon (32%), splice site (17%), and stop loss (4%) mutations (Figure 2C). These results show that a large number of high impact mutations (>1K) arise as the cannabis mother plant grows and is maintained for a long time and as a consequence potentially severely affects the function of important genes. Owing to the lack of any significant enrichment in terms of gene ontology (GO) annotation, we investigated the functional impact of these mutations in the important cannabinoid and terpene production related genes individually using public databases.

**Figure 2.**
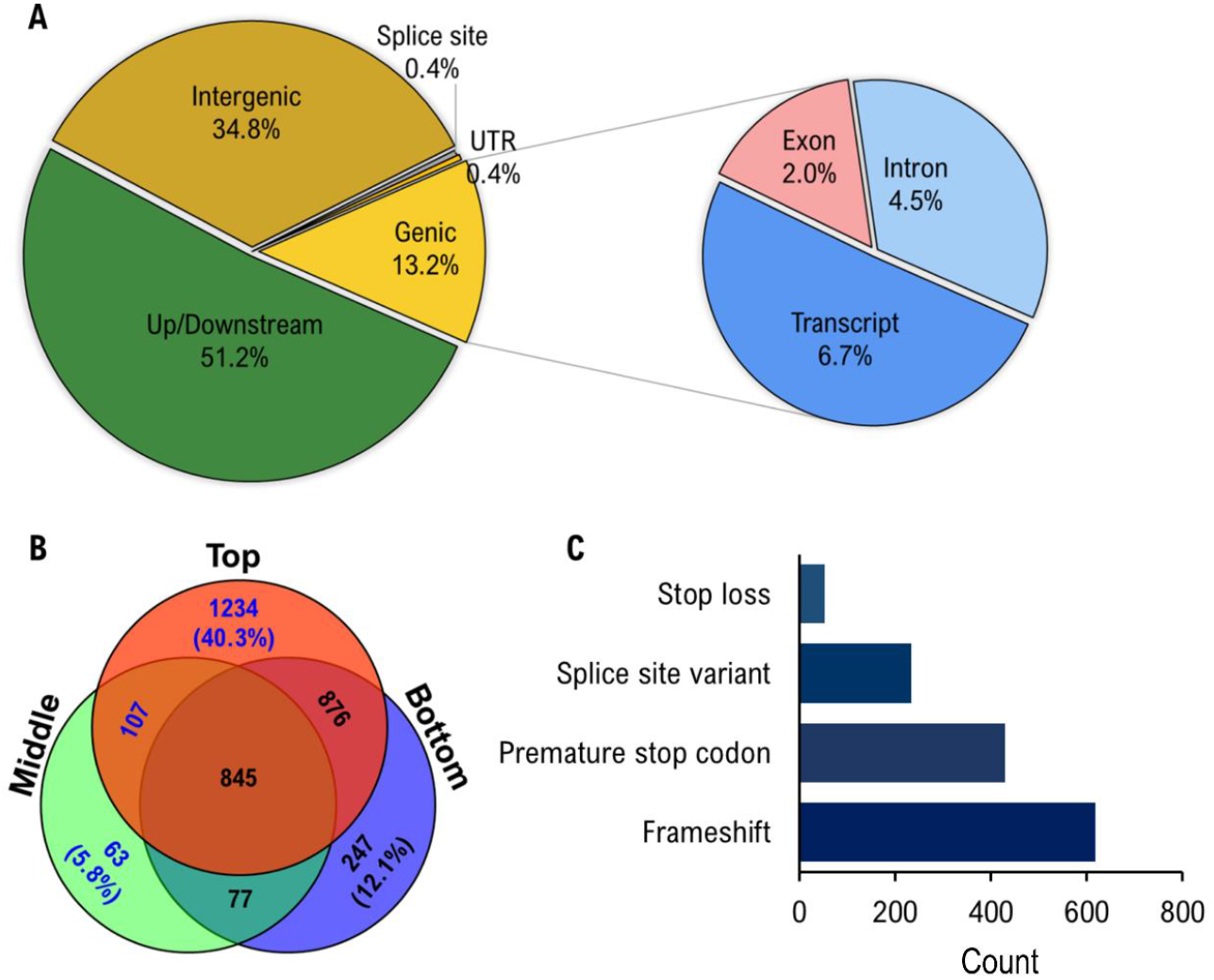
Analysis of the location and impact of nucleotide variants. (**A**) A pie graph visualization that displays the percentages from the five main categories of found variants (intergenic, up/downstream, genic, UTR, and splice site) while a secondary pie graph reveals the breakdown of the genic category (transcript, exon and intron). (**B**) A Venn diagram showing the high impact variants throughout and between the top, middle and bottom with the blue text highlighting *de novo* variants. (**C**) High impact mutations were organized into four categories and represented frameshift (46%), premature stop codon (32%), splice site (17%), and stop loss (4%) mutations.

### Properties of novel mutations on cannabinoid and terpene pathways genes

To more specifically focus on the potential impact of these mutations on secondary metabolite production, we studied the functional impact of mutations that occurred in or near the necessary cannabinoid and terpene pathway genes. We determined which enzymes were required to create essential chemical compounds and which chromosome they can be found on the cs10 v. 2.0 reference genome (Grassa *et al*., 2021) for both cannabinoid and terpene pathways from public databases (Table 2). As seen from the Table 2, we calculated and predicted the number and type of mutations, nucleotide diversity (π) in 20kb window encompassing these genes and the contrast of the π in these genic regions compared to the average genome wide. Analysis of the prediction of the functional impact of the mutations determined that none of these genes contained a high impact mutation and recorded all observed variants as modifier. The ratio of gene to genome wide π allowed us to provide a sense of potentially conserve or somatic mutation prone genes. The most notable genes that undergo somatic mutation were ones with ratio values greater than 2.0 which includes OLS, CBDAS, HMGR2 and CsTPS9FN, with values of 2.08, 4.35, 2.95, and 2.50, respectively. Of the remaining 40 genes, 32 were under 2.0 and the other eight are missing data due to no data found on the NCBI’s cannabis protein table. Overall, four genes are noteworthy because the number of variants emerging, and seem to be more prone to somatic mutations than others, but this will require additional research to determine if this is a common trend.

**Table 2.**
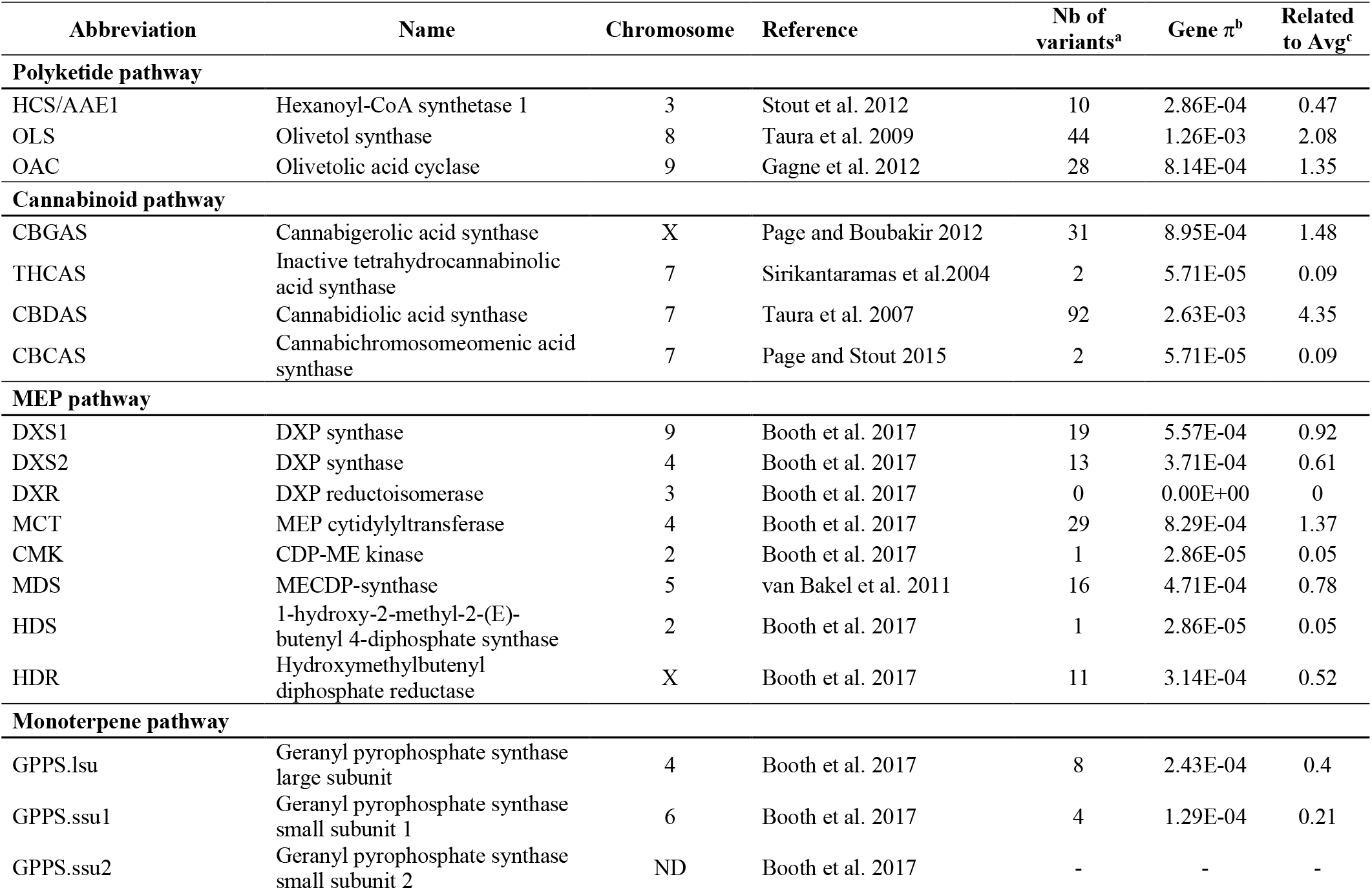
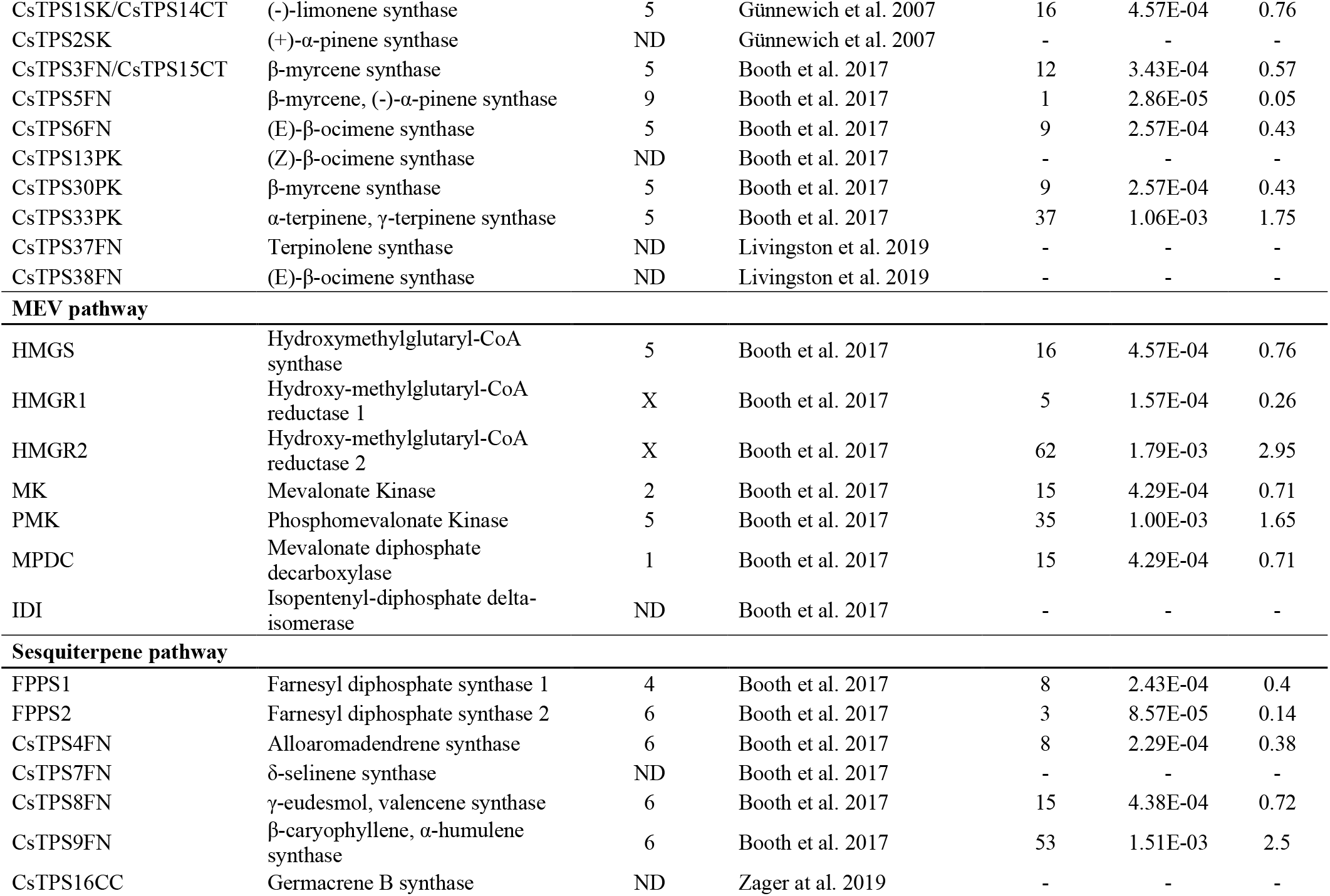
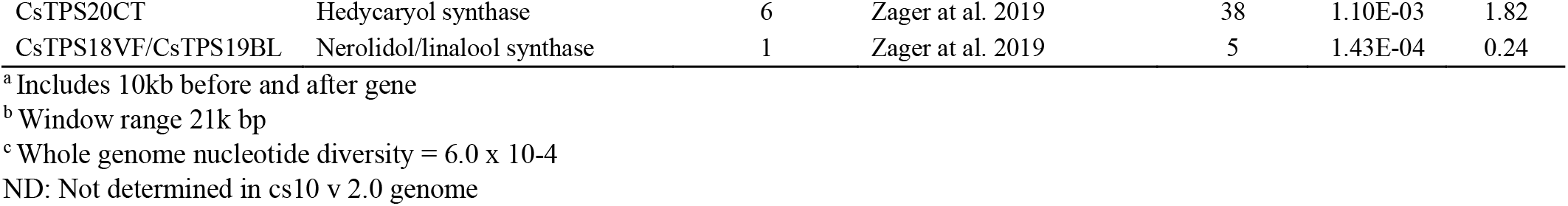
Variants in cannabinoid and terpene pathway genes

## Discussion

### Genomic diversity within cannabis

Numerous studies have investigated intra-plant somatic mutations in long-lived perennials, but few have examined this in annuals and this research is the first to look at intra-plant mutations in cannabis. (Diwan *et al*., 2014; Plomion *et al*., 2018; Hanlon, Otto and Aitken, 2019; Orr *et al*., 2020). Many previous studies used molecular markers to investigate and calculate mutation rates, which are prone to miss more rare mutations that WGS will capture. Some research on intra-plant genetic variation were unreliable as they used extrapolations from microsatellites or chlorophyll mutations, however, more recent research has been able uncover a more complete understanding due to advances and technologies becoming more accessible (Schoen and Schultz, 2019). For example, two studies using WGS on a long-lived Oak tree (*Quercus robur*) reported a much lower mutation rate than what was obtained from scaling up estimates from *Arabidopsis thaliana*. Based on this they hypothesized that perennials may have a mechanism to reduce the accumulation of mutations that is lacking in short lives annual species (Schmid-Siegert *et al*., 2017; Plomion *et al*., 2018). Likewise in mammals, it has been shown that mice have more mutations per cell division than humans, which is consistent with the hypothesis that short lived species have a higher mutation rates (Milholland *et al*., 2017). Currently, somatic mutations are believed to be common in plants but the mutation rate, distribution, morphological effects, age or size influence, and the differences between annuals and perennials remains poorly understood (Schoen and Schultz, 2019).

Naturally, cannabis is an annual species where it lives until its flowers are pollinated and seeds are produced. This all occurs during a single season that can range from a few months to closer to a year, and then it naturally dies. In contrast, cannabis plants maintained as mothers are artificially maintained in a perpetual vegetative state and replaced periodically using clonal propagaules that can extend their life span to several years or even decades. Based on Muller’s ratchet, a decline in plant vigour is likely during this period due to the accumulation of deleterious mutations and the absence of recombination. The extended life-span of mother plants significantly increases this concern because each *de novo* deleterious variant bears the potential for both multiplicative and cumulative effects to genome instability, altered gene expression, molecular heterogeneity, tissue disintegration and vulnerability to stress (Govindaraju *et al*., 2020). In other species, propagules that inherit deleterious mutation from a mother plant exhibited smaller leaves, reduced nutrient translocation capacity, degraded genetics, less rooting, lower plant vigour, and a decrease in growth (Wendling *et al*., 2014; Singh *et al*., 2015; Orr *et al*., 2020). Therefore, our results lead us to hypothesize that prolonging the lifespan of Cannabis plants and constantly pruning clonal cuttings is leading to the observed plant decline.

In this study, we called over 2 million nucleotide variants within a single cannabis plant. In comparison, the long-lived oak study only called 5,330 potential SNVs that had accumulated over many years (Schmid-Siegert *et al*., 2017). Interestingly, the Napoleon Oak genome was less than 1% heterozygous while the cannabis genome is known to be highly heterozygous (estimated at 12.5 – 40.5%) and contains substantial amounts of repetitive elements (estimated at 70%) (Hurgobin *et al*., 2020). We identified somatic mutations with deep sequencing (>50x depth of coverage) and used systematic filtering steps to reduce errors that may have occurred from next-generation sequencing (NGS) and incorrect mapping (Ajay *et al*., 2011). Additionally, to assess the possibility of inaccuracies, resequencing was performed on the original top tissue using the same protocol and procedure. The comparison between these samples were used to confirm the quality of genotyping and revealed an astonishing 99.97% agreement in variant calling. Thus, indicating that errors or incorrect mapping had nearly zero effect on the callings of variants.

A recent cannabis study sequenced 40 cannabis genomes and they reported an average of 12.8 million SNPs+Indels for dispensary grade cannabis (Type I and Type II plants) which equated to a variant every ~73 bases while our results equated to a variants every ~425 bases (McKernan *et al*., 2020). Although our variants were six-fold less than the average found in the other study, partly due to clonal origin of our samples, they still represent a substantial quantity of variants that contain the ability to interfere with the stability and quality of the plant.

One might assume that calculating the ratio of transition to transversion from three samples derived from one individual mother plant is less interesting from a statistical point of view, but the comparison of this ratio with other cannabis research and other plant species was appealing. The Ts/Tv ratio found in this study (1.88) was similar to a recent study (Soorni *et al*., 2017) where the average Ts/Tv ratio for 69 cannabis individuals was 1.65, showing an intriguing level of intra-plant genomic diversity. As the Ts/Tv ratio is comparable, we can verify that the value is within an expected range, which reduced the chances of high false positives or bias. Compared to other plant species, this ratio is similar to oil palm (1.67) and significantly higher than maize (1.02) (Batley *et al*., 2003; Pootakham *et al*., 2015).

### Variants across intra-plant genome

We expected to witness a systematically hierarchical nature of mutation accumulation from the bottom to the top, as was seen by Diwan *et al* (2014) in their research on genome differences within Yoshino cherry (*Prunus* × *yedoensis*) and Japanese beech (*Fagus crenata*) trees. In this study, we witnessed a minor drop in total variants between the bottom and middle compared to the middle to top. Although we didn’t observe a uniform increase, the uppermost sample was the more genetically distant from the bottom than was the middle, as seen by other previous intra-plant studies (Schmid-Siegert *et al*., 2017; Plomion *et al*., 2018; Hanlon, Otto and Aitken, 2019). Additionally, the primary analytical focus in this study was the middle and top variants because they represent *de novo* mutations while the bottom section corresponds to the oldest growth tissue where, in theory, the mutations counts should be the lowest.

While many uncertainties remain about somatic mutation in plants, our results demonstrating significant genetic variation may be explained by a few possibilities 1) environmental factors 2) long-term pruning from the top 3) difference between perennials and annuals 4) small sample size. Environmental factors have been well studied and are known to impact mutation rates, especially during shock or stress (Gill *et al*., 1995). Thus, any changes to the environment during the growth and maintenance of the mother plant, such as light or temperature stress, may have altered the mutation rates and contributed to their accumulation over time. Next, due to the nature of cannabis mother plants, they are consistently apically pruned and clonal propagules are taken for propagation. This practice has a direct impact on the balance between auxin and cytokinin levels, promotes rapid growth from originally inhibited axillary bud sites, and as a result could impact the rate of mutations within the plant (Prusinkiewicz *et al*., 2009; Schaller *et al*., 2015). Recently, NGS studies identified major differences in the rate of mutations between perennials and annuals species, with perennials having a significantly lower rate than expected, suggesting they may possess a mechanism to suppress the rate (Schoen and Schultz, 2019). These findings are consistent with our results that found a relatively high number of mutations and lead us to believe that annuals contain a more severe and sporadic mutation rate such that prolonging their normal lifespan may lead to a substantial mutation load with potentially deleterious effects. Lastly, due to the expensive costs of full genome sequencing and intention to simply identify intra-plant variations, only three positions (i.e. bottom, middle, and top) were sequenced with the ability to call ultra-rare mutations. As such, it is unknown if the degree of diversity observed in this study is representative of the species. Further work is needed using larger sample sizes across multiple genotypes in different environments to gain a better understanding of this phenomenon.

### Impact on cannabinoid and terpene synthase

We investigated the impact that these mutations may have had on both the cannabinoid and terpene pathways because of their medical and product quality importance (Hanuš and Hod, 2020; Singh *et al*., 2020). First, we examined mutations that were categorized as high impact from analysis using SnpEff (Cingolani *et al*., 2012). These mutations are known to have a large impact on protein production, protein truncation, loss of function or trigger nonsense mediated decay (Cingolani *et al*., 2012). Although we didn’t identify any high impact mutations in the cannabinoid and terpene synthase genes, the analysis revealed a similar arrangement of the distribution of mutations from the total number of variants. Interestingly, the top sample contained an even larger total percentage by ~6% while the middle had 3% less and the bottom had a less than 1% difference. This further supported of the notion that apical regions are more genetically distant than the basal regions.

Furthermore, we examined mutations in these pathway genes that fell under the category of *moderate*, *low*, or *modifier* which relates to non-disruptive changes to protein effectiveness, mostly harmless or unlikely to change protein performance, and non-coding variants where predictions are difficult or there is no evidence of impact, respectively. Our analysis revealed that all discovered variants fell under the modifier category and encouraged the assessment of nucleotide diversity (π) with a 20kb scope of critical genes for both pathways. Most of the 44 genes investigated had a similar or lower value than the average nucleotide diversity (π), however, four major genes had values over double. Two of these are a part of terpene production (HMGR2 and CsTPS9FN) and the other two are involved in the cannabinoid pathway (OLS and CBDAS). HMGR2 role is to convert acetyl-CoA into mevalonic acid which undergoes a few more steps to produce the synthesis of both γ-eudesmol or β-caryophyllene terpenes. CsTPS9FN is the last enzyme necessary to produce the β-caryophyllene terpene. OLS and CBDAS were particularly intriguing because OLS is an essential enzyme for the whole cannabinoid pathway as it converts hexanoyl-CoA into olivetolic acid which then converts into CBGA, a precursor to many well-known cannabinoids (i.e. THCA, CBDA, CBCA). CBDAS is the last enzyme required to convert CBGA into CBDA and this is especially important as its the main cannabinoid used to treat various health concerns (Maroon and Bost, 2018). Both critical enzymes revealed a more than double nucleotide diversity (π) above average and as a result could indicate initial signs of decay in the cannabinoid pathway.

Mother plants are usually selected through large scale, costly, screening programs, and marketed as strains with unique properties. The alteration of genes over time represents a significant challenge in long-term batch to batch consistency. Also, cannabis used for medicinal purposes must ensure a consistent product that provides the appropriate properties and quality that are necessary for treatments. Thus, a greater insight into somatic mutations may enable new or superior procedures to assist the preservation of elite cultivars in clonally propagated plants. Overall, our research highlights an important phenomenon related to maintaining elite genetics and could provide an underlying mechanism for the decay of cannabinoids and plant vigour that has been anecdotally observed (Cannabis growers, personal communication, 2020).

## Conclusion

The findings in this study demonstrate that the genetic diversity exists with a single cannabis plant and the genetic mosaicism hypothesis applies to *C. sativa*. This study is the first to investigate the existence of this phenomenon in cannabis plants and the potential consequences from accumulating somatic mutations in an artificially prolonged annual species. As cannabis normally lives for ~3-6 months, this process likely enables an unknown, but manageable, amount of somatic mutations to accumulate. Currently, somatic mutations in plants have many uncertainties remaining, but due to modern genetic technologies and more affordable WGS, there has been more contributions with higher degrees of accuracy and precision on this topic. From a practical standpoint, this significantly benefits the cannabis industry as understanding this phenomenon will help establish best practices for maintaining mother plants to minimize, slow or prevent the accumulation of mutations. Based on these data, we advocate replacing mother plants using cuttings from the basal portion of the plant and discourage excessively extending the life of a mother plant. Additionally, important genetics should be preserved using cryopreservation techniques where the original genetic profile can be maintained and accessed indefinitely (Uchendu *et al*., 2019). The research here provides a concrete basis for cannabis mutation research. However, the current study lacked different cultivars, generational data, mutation rates and multiple biological replicates. Thus, future research will be necessary to enhance and solidify our understanding of somatic mutations and the mutagenic potential that exists within a cannabis mother plant.

## Materials and Methods

### Plant material

A high-THCA mother plant of *Cannabis sativa* cv. “Honey Banana” (15-20% THC; <1% CBD), was grown indoors at BrantMed Inc. in Ontario, Canada. The plant was grown in nutrient rich soil with regular feeding of a full nutrient solution developed for vegetative growth adjusted to ~6.5 pH. The proprietary nutrient solution was relatively high in nitrogen, with moderate levels of phosphorous, potassium (NPK) and micronutrients, administered every 3-5 days as needed. The plant was grown from seed and transplanted to larger pots as needed, until reaching the 76L (20gal) pot where it remained until this study was conducted. The environmental conditions were maintained at 20-25°C and 55-65% relative humidity using BrantMed Inc LED lighting (Grow Light E1-300W) under long day photoperiods at a 18:6 hour light:dark cycle to maintain the mother plant in an indefinitely vegetative state. These broad “white” spectrum lights (4.2% red 650-670 nm) with 300-watts provided a photosynthesis photon flux of >440 μmol/s and came with a PAR photon efficacy of 2.2 μmol/J. Samples were removed when the plant had reached an age of approximately 1.5 years. We isolated ~2.5cm of fresh stem tissue from three location at 59cm, 151cm, and 226cm which represented the bottom, middle and top samples, respectively (Figure 1A). The samples were frozen and stored in a freezer until DNA extraction.

### DNA extraction and whole genome resequencing

Frozen stem tissues were ground using a Qiagen TissueLyser. DNA was extracted from approximately 100 mg of ground tissue using the Qiagen Plant DNeasy Mini Kit according to the manufacturer’s protocol. DNA was quantified on a NanoDrop spectrophotometer and on a Qubit fluorometer. Illumina Paired-End libraries were constructed for three DNA samples using the Illumina Tru-seq DNA Library Prep Kit (Illumina, San Diego CA, USA) following the manufacturer’s instructions. The quality of DNA library was verified on an Agilent Bioanalyzer with a High Sensitivity DNA chip. The sequencing was performed on an Illumina NovaSeq 6000 platform at the McGill University-Génome Québec Innovation Center in Montreal, QC, Canada generating >1 billion 150-bp paired-end reads to provide >50x death of coverage by sequencing reads against the mapped the public domain *C. sativa* reference genome (Grassa *et al*., 2021). A second whole-genome resequencing was carried out using similar protocols and procedures as previously mentioned onto the original frozen tissue used for the top sample. The Illumina NovaSeq generated 172M reads with a depth of coverage of 32x.

### Bioinformatic data analysis

Illumina paired-end reads were processed using Fast-WGS bioinformatics pipeline (Torkamaneh *et al*., 2018). In summary, the reads were mapped against the cs10 v 2.0 cannabis reference genome (GenBank Accession No. GCA_900626175.2; Grassa *et al*., 2021) using BWA-MEM (Li, 2013). This cannabis reference was chooses as it is currently the most complete reference and has been proposed to be the cannabis reference for genomics by the International Cannabis Genomics Research Consortium (ICGRC; Hurgobin *et al*., 2020). The nucleotide variants were called using Platypus (Rimmer *et al*., 2014). In general, we removed variants if: 1) they had more than two alleles, 2) an allele was not supported by reads on both strands, 3) the overall quality (QUAL) score was <32, 4) the mapping quality (MQ) score was <20, 5) read depth (minNR) was <10 and 6) the number of reads supporting variant (minNV) was <10. Functional annotation of nucleotide variation was performed by SnpEFF and SnpSift (Cingolani *et al*., 2012) using a customized reference built using cannabis reference genome annotation file downloaded from NCBI. To obtain the description of genes with large impact, we used the NCBI’s protein table for cannabis sativa database. The gene ontology (GO) analysis was done using the singular enrichment analysis (SEA) method implemented in agri-GO (Du *et al*., 2010).

### Diversity, LD and clustering analysis

Nucleotide diversity (π) was calculated using VCFtools (Danecek *et al*., 2011), with a window of 20K bp on the full dataset. An average π across all windowed calculations was used to obtain a genome-wide average π. A neighbor-joining phylogenetic tree was constructed using full dataset in Tassel 5.0 (Bradbury *et al*., 2007). The linkage disequilibrium (LD) decay was determined using PopLDdecay version 3.40 Beta (Zhang *et al*., 2019). IGV 2.8 (Robinson *et al*., 2011) was utilized to display the distribution of variants within the three samples which was produced with the indexed version of the VCF file.

## Accession number

The accession number is not currently available but will be able from NCBI before official publication.

## Acknowledgments

This work was conducted as part of a collaborative research project funded by Brantmed Inc. and NSERC Alliance [#ALLRP555969 −20 to A.M.P.J and D.T.]. The authors wish to thank BrantMed Inc. for supporting this project.

## Author contributions

DT, AMPJ, and KA conceived the project. DT carried out the WGS and variant calling. KA performed bioinformatics analysis. KA, DT, and AMPJ contributed to writing the manuscript.

## Supplementary materials

**Figure S1**. Comparison of nucleotide variants from two separate WGS of the top sample.

**Figure S2.** LD decay graph revealing a rapid decay in only a few kb.

**Figure S3.** Visual representation of nucleotide variants distributed throughout the chromosomes.

