## Supplemental Data for "Accumulation of somatic mutations leads to genetic mosaicism in Cannabis"

### Supplemental Materials

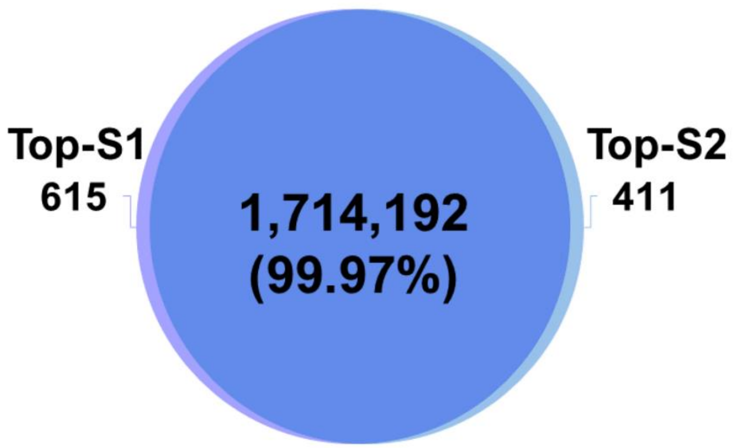


**Figure S1.** Comparison of nucleotide variants called from the first and second runs of WGS on the original top tissue with a near identical agreement in variant calling.

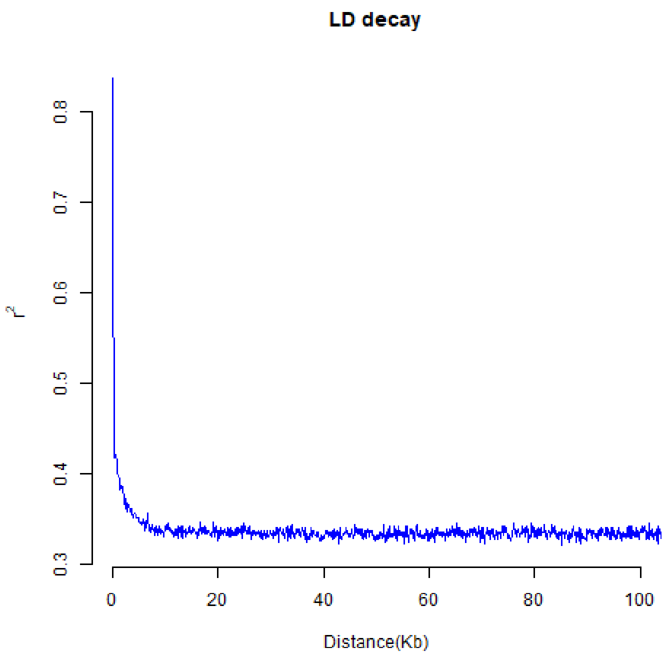


**Figure S2.** LD decay graph that shows a rapid decay to half in merely a few kb.

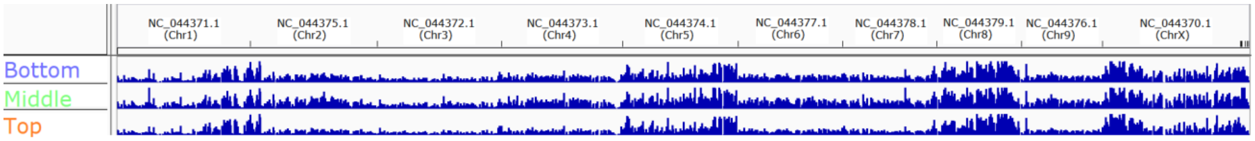


**Figure S3.** A visual representing the distribution of nucleotide diversity through the full genome from the bottom, middle and top samples. Chromosomes 1, 5, 8 and X are under a heavier burden of nucleotide variants.
